# The EVcouplings Python framework for coevolutionary sequence analysis

**DOI:** 10.1101/326918

**Authors:** Thomas A. Hopf, Anna G. Green, Benjamin Schubert, Sophia Mersmann, Charlotta P. I. Schäerfe, John B. Ingraham, Agnes Toth-Petroczy, Kelly Brock, Adam Riesselman, Chan Kang, Christian Dallago, Chris Sander, Debora S. Marks

**Affiliations:** Department of Systems Biology, Harvard Medical School, Boston, MA 02115, USA; Department of Cell Biology, Harvard Medical School, Boston, MA 02115, USA; cBio Center, Department of Biostatistics and Computational Biology, Dana-Farber Cancer Institute, Boston, MA 02215, USA; Center for Bioinformatics, University of Tübingen, 72076 Tübingen, Germany; Applied Bioinformatics, Dept. of Computer Science, 72076 Tübingen, Germany; Max Plank Institute of Molecular Cell Biology and Genetics, 01307 Dresden, Germany; Program in Biomedical Informatics, Harvard Medical School; Department of Informatics, Technische Universität München, 85748 Garching, Germany

## Abstract

**Summary:** Coevolutionary sequence analysis has become a commonly used technique for de novo prediction of the structure and function of proteins, RNA, and protein complexes. This approach requires extensive computational pipelines that integrate multiple tools, databases, and data processing steps. We present the EVcouplings framework, a fully integrated open-source application and Python package for coevolutionary analysis. The framework enables generation of sequence alignments, calculation and evaluation of evolutionary couplings (ECs), and de novo prediction of structure and mutation effects. The application has an easy to use command line interface to run workflows with user control over all analysis parameters, while the underlying modular Python package allows interactive data analysis and rapid development of new workflows. Through this multi-layered approach, the EVcouplings framework makes the full power of coevolutionary analyses available to entry-level and advanced users.

**Availability:** https://github.com/debbiemarkslab/evcouplings

## 1 Introduction

Coevolutionary sequence analysis presents a promising new approach to the long-standing problem of *de novo* prediction of the 3D structure of proteins and RNAs. Pairwise graphical models of biological sequences are used to identify evolutionary couplings (ECs) between sites, which frequently correspond to physical contacts between nucleotides or amino acids in the molecule’s three-dimensional structure. ECs have been used successfully to predict the tertiary contacts (Marks, et al., 2011; Morcos, et al., 2011) and full 3D structure of proteins (Marks, et al., 2011) and RNA (Weinreb, et al., 2016), as well as protein-protein and protein-RNA interactions, disorder and conformational changes (Weigt, et al., 2009; Hopf, et al., 2012; Hopf, et al., 2014; Ovchinnikov, et al., 2014; Rodriguez-Rivas, et al., 2016; Toth-Petroczy, et al., 2016) and the effects of mutations (Figliuzzi, et al., 2015; Hopf, et al., 2017). More recently, hybrid approaches have integrated ECs with experimental data for improved structure determination using NMR, cryo-EM and X-ray crystallography data (Tang, et al., 2015; Sjodt, et al., 2018).

**Figure 1:**
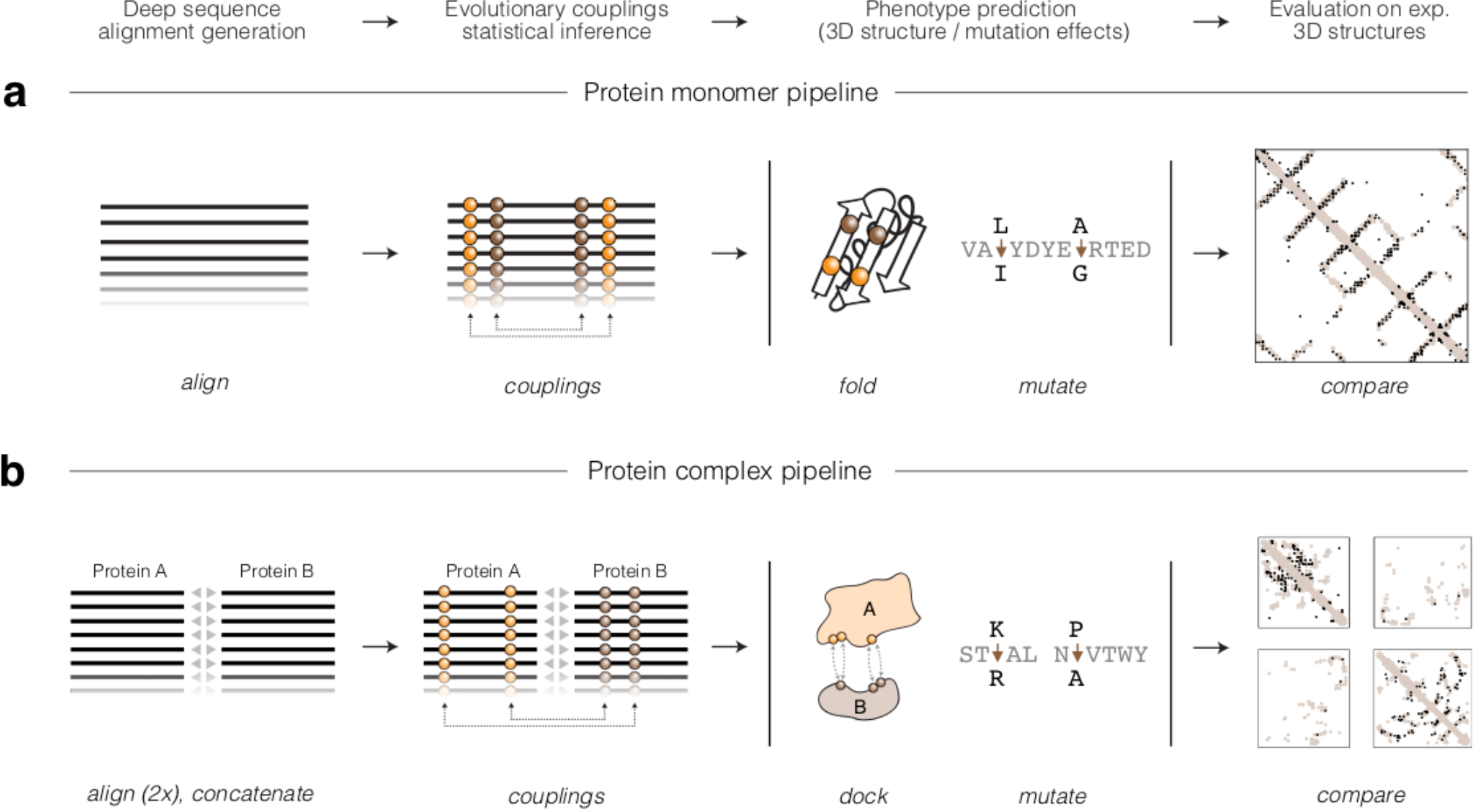
The EVcouplings Python framework. (a) Protein monomer EVcouplings pipeline. A multiple sequence alignment (MSA) of a query protein is generated (align stage) by searching a sequence database. Given the MSA, a pairwise undirected graphical model is inferred to identify evolutionary coupled positions (ECs, couplings stage). Optionally, the identified EC and model parameters can be used as constraints to fold the protein (fold stage), predict mutational effects (mutate stage), and compare the predicted contacts to experimental structures (compare stage). (b) Protein complex EVcouplings pipeline. Two monomer alignments are generated independently (align stage), and then a joined alignment is constructed by pairing putatively interacting partners (concatenate stage). A pairwise undirected graphical model is inferred to identify inter-and intra-molecular ECs (couplings stage), which then can be used as constraints to dock (dock stage), predict mutational effects (mutate stage), or compare the predicted tertiary structure with experimental structures of monomers or complexes (compare stage).

However, all of these applications require complex analysis pipelines using multiple tools and data sources, as well as extensive data pre-and post-processing. Most software development in the field has focused on high-performance reimplementations of EC inference tools (Kaján et al., 2014; Seemayer, et al., 2014), and one addressed format conversion between different alignment and inference tools (Simkovic, et al., 2017). To make EC methods accessible to a general biological audience, following an earlier preliminary implementation (Sheridan, et al., 2015), we present the first open source application and Python package for end-to-end evolutionary coupling analysis. EVcouplings covers all necessary functionality, including alignment generation, EC calculation, *de novo* structure and mutation effect prediction, visualization of results, and comparison of predictions to experimental structures.

## 2 EVcouplings framework

The EVcouplings framework integrates the functionality of the previously published methods EVfold (Marks, et al., 2011; Hopf, et al., 2012), EVcomplex (Hopf, et al., 2014), and EVmutation (Hopf, et al., 2017). It provides (1) an easy-to-use command-line application and (2) a modular Python package containing all functions, data structures, and pipelines that comprise the application.

### 2.1 Command-line application

The command-line application allows users to obtain predictions for their proteins and complexes of interest by running the respective EVcouplings pipelines. Each pipeline is comprised of a series of modular stages that can be configured using a YAML file, aiding reproducibility by documenting all parameters. The pipeline can be halted and restarted at any stage, allowing the inspection of intermediate results and partial reruns to explore the effects of different parameters. The pipeline is parallelized and supports local multi-process execution as well as commonly used cluster systems, and automatically handles job submission and monitoring.

The major steps of the prediction pipelines are as follows (pipeline stages in brackets):

i. **Alignment generation *(align, concatenate)*:** The key prerequisite to infer evolutionary couplings is a high-quality sequence alignment of sufficient evolutionary depth. EVcouplings supports the generation of alignments using jackhmmer and hmmsearch (Eddy, 2011), as well as the import of externally computed alignments. Since the identification of an optimal evolutionary depth can be challenging, the application supports the parallel exploration of different sequence inclusion thresholds, provides flexible alignment filtering parameters, and calculates summary statistics to assess alignment quality. When predicting protein complexes, the alignment generation of individual proteins is followed by matching putatively interacting proteins to create a joint alignment of the complex (*concatenate* stage), using one of several provided approaches including genomic co-location and homology identification with bidirectional best hits.
ii. **EC inference *(couplings)*:** ECs are then inferred from the alignment using a pairwise undirected graphical model of sequences. The EVcouplings framework offers the two major established inference methods: (a) pseudolikelihood maximization (Balakrishnan, et al., 2011; Ekeberg, et al., 2013), using plmc (https://github.com/debbiemarkslab/plmc), and (b) a Python reimplementation of the mean-field method (Marks, et al., 2011). ECs are scored probabilistically with a two-component mixture model to identify high confidence couplings (Toth-Petroczy, et al., 2016).
iii. **3D structure prediction *(fold)*:** The 3D structure of proteins is predicted with the molecular dynamics software CNS (Brunger, 2007) using ECs as distance restraints on residue pairs in an extended polypeptide. Models are clustered and ranked according to geometric criteria and evaluated against experimental structures if available (Marks, et al., 2011).
iv. **Mutation effect prediction *(mutate)*:** Based on the inferred probability model of sequences, mutational landscapes and the effects of user-provided mutations are computed and visualized, allowing the user to predict the phenotypic impact of genetic variation (Hopf, et al., 2017)
v. **Comparison to experimental structures *(compare)*:** Where experimental structures of the target protein can be identified, the agreement between ECs and residue contacts in 3D is evaluated and visualized using contact maps (2D residue-residue contact plots) and PyMOL scripts (Schrodinger, 2015). EVcouplings offers comprehensive support for finding homologs in the PDB using sequence searches, index mapping between target sequence and residues in the structure, and identification of intra-protein, inter-protein and homomultimeric residue contacts.

### 2.2 EVcouplings Python Package

The command-line application is built on the underlying *evcouplings* Python package, whose modular architecture and comprehensive documentation facilitate the development of new stages and pipelines. Additionally, the package serves as a toolbox for handling and analyzing EC-related data. For example, the *CouplingsModel* class allows the user to read and explore graphical model parameters and to perform mutation effect calculations on sequence variants. The submodule *evcouplings*.*compare* provides an integrated solution to aligning and handling 3D structure data based on the MMTF PDB format (Bradley, et al., 2017) and SIFTS mappings (Velankar, et al., 2013). Examples for interactive usage are provided in Jupyter notebooks (Kluyver, et al., 2016) distributed with the package, and extensive documentation is available on the web (http://evcouplings.readthedocs.io).

## 3 Conclusion

EVcouplings is the first open source, integrated pipeline for evolutionary couplings analyses. The underlying API serves as a modular basis for data analysis and further developments and will allow developers to create new workflows as the field advances.

*Conflict of Interest:* none declared.

## References

Balakrishnan, S., et al. Learning generative models for protein fold families. Proteins 2011;79(4):1061–1078.

Bradley, A.R., et al. MMTF-An efficient file format for the transmission, visualization, and analysis of macromolecular structures. PLoS Comput Biol 2017;13(6):e1005575.

Brunger, A.T. Version 1.2 of the Crystallography and NMR system. Nature protocols 2007;2(11):2728.

Eddy, S.R. Accelerated profile HMM searches. PLoS computational biology 2011;7(10):e1002195.

Ekeberg, M., et al. Improved contact prediction in proteins: using pseudolikelihoods to infer Potts models. Phys Rev E Stat Nonlin Soft Matter Phys 2013;87(1):012707.

Figliuzzi, M., et al. Coevolutionary landscape inference and the context-dependence of mutations in beta-lactamase TEM-1. Molecular biology and evolution 2015;33(1):268–280.

Hopf, T.A., et al. Three-dimensional structures of membrane proteins from genomic sequencing. Cell 2012;149(7):1607–1621.

Hopf, T.A., et al. Mutation effects predicted from sequence co-variation. Nature biotechnology 2017;35(2):128.

Hopf, T.A., et al. Sequence co-evolution gives 3D contacts and structures of protein complexes. Elife 2014;3.

Kaján, L., et al. FreeContact: fast and free software for protein contact prediction from residue co-evolution. BMC Bioinformatics 2014;15(1):85.

Kluyver, T., et al. Jupyter Notebooks-a publishing format for reproducible computational workflows. In, ELPUB. 2016. p. 87–90.

Marks, D.S., et al. Protein 3D structure computed from evolutionary sequence variation. PloS one 2011; 6(12):e28766.

Morcos, F. et al. Direct-coupling analysis of residue coevolution captures native contacts across many protein families. Proc Natl Acad Sci USA 2011;108(49):E1293–1301.

Ovchinnikov, S., Kamisetty, H. and Baker, D. Robust and accurate prediction of residue-residue interactions across protein interfaces using evolutionary information. Elife 2014;3.

Rodriguez-Rivas, J. et al. Conservation of coevolving protein interfaces bridges prokaryote-eukaryote homologies in the twilight zone. Proceedings of the National Academy of Sciences 2016;113(52): 15018–15023.

Schrodinger, LLC. The PyMOL Molecular Graphics System, Version 1.8. In.; 2015.

Seemayer, S., Gruber, M. and Soding, J. CCMpred—fast and precise prediction of protein residue-residue contacts from correlated mutations. Bioinformatics 2014;30(21):3128–3130.

Sheridan, R. et al. EVfold. org: Evolutionary Couplings and Protein 3D Structure Prediction, biorxiv 2015:021022.

Simkovic, F., Thomas, J.M.H. and Rigden, D.J. ConKit: a python interface to contact predictions. Bioinformatics 2017;33(14):2209–2211.

Sjodt, M. et al. Structure of the peptidoglycan polymerase RodA resolved by evolutionary coupling analysis. Nature 2018.

Tang, Y. et al. Protein structure determination by combining sparse NMR data with evolutionary couplings. Nature methods 2015;12(8):751.

Toth-Petroczy, A., et al. Structured states of disordered proteins from genomic sequences. Cell 2016;167(1):158–170. el12.

Velankar, S., et al. SIFTS: Structure Integration with Function, Taxonomy and Sequences resource. Nucleic Acids Res 2013; 41(Database issue):D483–489.

Weigt, M. et al. Identification of direct residue contacts in protein-protein interaction by message passing. Proceedings of the National Academy of Sciences 2009;106(1):67–72.

Weinreb, C. et al. 3D RNA and functional interactions from evolutionary couplings. Cell 2016;165(4):963–975.

